# LKB1 acts as a critical brake for the glucagon-mediated fasting response

**DOI:** 10.1101/2022.01.29.478228

**Authors:** Suehelay Acevedo-Acevedo, Megan L. Stefkovich, Sun Woo Sophie Kang, Rory P. Cunningham, Constance M. Cultraro, Natalie Porat-Shliom

## Abstract

As important as the fasting response is for survival, an inability to shut it down once nutrients become available can lead to exacerbated disease and severe wasting. The liver is central to transitions between feeding and fasting states, with glucagon being a key initiator of the hepatic fasting response. However, the precise mechanisms controlling fasting are not well defined. One potential mediator of these transitions is Liver Kinase B1 (LKB1) given its role in nutrient sensing. Here, we show LKB1 knockout mice have a severe wasting and prolonged fasting phenotype despite increased food intake. By applying RNA sequencing and intravital microscopy we show that loss of LKB1 leads to a dramatic reprogramming of the hepatic lobule through robust upregulation of periportal genes and functions. This is likely mediated through the opposing effect LKB1 has on glucagon pathways and gene expression. Conclusion: our findings show that LKB1 acts as a brake to the glucagon-mediated fasting response resulting in “periportalization” of the hepatic lobule and whole-body metabolic inefficiency. These findings reveal a new mechanism by which hepatic metabolic compartmentalization is regulated by nutrient-sensing.

## Introduction

Fasting is a conserved, adaptive response to nutrient deprivation that involves metabolic and transcriptional changes. The liver is critical in this response by sequentially activating catabolic and inhibiting anabolic processes to meet energy demands. Glucagon is at the crux of regulating the hepatic metabolic adaptation to fasting. When circulating glucose levels drop, glucagon stimulates hepatic glucose output via its release from glycogen storages or *de novo* synthesis through gluconeogenesis, as well as fatty acid oxidation and ketogenesis which provide crucial fuels to peripheral tissues during extended fasting ^1^. Equally important is the ability to shut down the fasting response once nutrients become available. This action prevents catabolic processes from continuing needlessly and preventing gluconeogenesis, lipolysis, and ketogenesis to go unchecked. Indeed, impairment of fasting inhibition is observed in Non-Alcoholic Fatty Liver Disease (NAFLD) patients where gluconeogenesis and lipolysis are elevated ^2, 3^. Indeed, metformin and glucagon-like peptide 1 (GLP-1) agonists have been successful in treatment of obesity, type 2 diabetes, and NAFLD via attenuation of the glucagon response ^4-7^. Fine tuning of this complex response is temporally regulated by glucagon through activation of transcription factor cascades leading to sequential waves of gene expression that optimize energy usage and prevent tissue damage ^8^.

Recently, *Cheng at al*. demonstrated that glucagon is also involved in the spatial regulation of hepatic metabolism ^9, 10^. The spatial segregation of hepatic metabolism, also known as liver zonation, permits simultaneous operation of conflicting pathways ^11-13^. This is achieved through the organization of hepatocytes in polarized, hexagonal lobules in which six portal vessels deliver oxygen, nutrients, and hormones (Sup Fig 1B) that flow directionally towards a single central vein. Several factors regulate zonation by affecting differential gene expression along the periportal (PP) to pericentral (PC) axis. For example, hypoxia and Wnt ligands activate β-catenin signaling resulting in the expression of target genes in PC cells ^14, 15^. On the other hand, glucagon controls gene expression in the PP zone ^10^. While the ‘push and pull’ relationship between Wnt ligands and glucagon have been established as a central determinant exerting zone-dependent expression patterns, it is not clear how hepatocytes integrate those signals during periods of fasting to maintain homeostasis.

One potential mediator is Liver kinase B1 (LKB1), given its role as a nutrient sensor that activates AMP-activated Protein Kinase (AMPK) and members of the AMPK-related (ARKs) family. LKB1 is also an important regulator of hepatic polarity ^16-19^, a feature that may be important for the spatial organization of the lobule. Loss of LKB1 in whole body ^20^ and liver-specific knockout ^18, 19, 21^ result in stark metabolic changes resembling cachexia, a condition that causes extreme weight loss due to fat and muscle wasting. Hepatic LKB1 was shown to be pivotal for regulating glucose homeostasis, as it is known to suppress gluconeogenesis through the activation of Salt-induced Kinase, ^22, 23^, and conversely, mice lacking LKB1 display elevated hepatic gluconeogenesis ^22^, driven by amino acid catabolism ^21^. Further, hepatic LKB1 is required for the glucose lowering effects of metformin ^22^. Despite its established involvement in whole-body homeostasis and gluconeogenesis, it is not clear how LKB1 loss of function results in such a severe metabolic phenotype.

Here, we show that hepatocyte-specific deletion of LKB1 (KO) leads to a prolonged fasting response in the liver resulting in a severe wasting phenotype and disrupted glucose homeostasis. At the transcriptional level, liver from KO mice displayed upregulated glucagon-induced genes throughout the hepatic lobule, many of which are enriched in PP regions in wild type (WT) mice. Our results suggest that LKB1 spatially fine-tunes glucagon pathways across the hepatic lobule to balance the extent of the fasting response which is critical for whole-body energy homeostasis. These findings further understanding of hepatic physiology and factors regulating liver zonation.

## Material and methods

### Reagents

Primary antibodies for western blots: LKB1 (3047S, 1:1000; Cell Signaling Technologies); β-tubulin (2146S, 1:1000; Cell Signaling Technologies); phosphoenolpyruvate carboxykinase 1 (PCK1 [ab28455], 1:1000, Abcam); Glyceraldehyde 3-phosphate dehydrogenase (GAPDH, 2118S, 1:1000; Cell Signaling Technologies). Secondary antibodies used: goat anti-mouse IgG, HRP conjugate (12-349, 1:1000, Millipore Sigma), goat anti-rabbit IgG, HRP conjugate (12-348, 1:1000, Millipore Sigma).

Primary antibodies used for immunofluorescence: Alexa Fluor 421, 647-conjugated CD324 (E-cadherin, 1:50, added during secondary antibody incubation, BioLegend); and Cytochrome P450 2E1 (Cyp2e1, ab28146, 1:200, Abcam). Secondary antibodies were purchased from Jackson ImmunoResearch. Alexa Fluor 647-phalloidin (1:200, Invitrogen) to label actin.

### Mice

The study was conducted according to the animal protocols approved by the National Cancer Institute (NCI)-Bethesda Animal Care and Use Committee (ACUC) protocol number LCMB037. C57BL/6J and LKB1 fl/fl mice (Jackson Laboratories) with free access to water and a standard diet (NIH-31 Open Formula, Envigo). Experiments were done during the light cycle.

### LKB1 KO

LKB1 fl/fl were injected with adeno-associated virus 8 (AAV8; 1.25 × 10^11^ GC) expressing Cre recombinase under the hepatocyte thyroid hormone-binding globulin (TBG) promoter (Vector Biolabs). AAV8 was delivered by tail vein injection to LKB1 fl/fl and C57BL/6J at 6 weeks of age and mice were analyzed 4 weeks after. As expected, male mice had higher body weights and lower fat mass than female mice (data not shown). However, since no sex by KO interaction was observed for any measurement, male and female data was combined.

### Liver lysate and western blots

Livers were snap-frozen in liquid nitrogen. Fifty miligram of tissue was homogenized in 3 mL RIPA buffer (Invitrogen) with 1X Halt protease and phosphatase inhibitor (Invitrogen). Homogenate was centrifuged at 13,000 rpm for 20 minutes at 4°C, and the supernatant was collected. Protein concentration was determined using a Pierce™ BCA Protein Assay Kit (Thermo Fisher Scientific). Lysates were boiled and 20-60μg of protein was loaded onto Mini-PROTEAN^®^ TGX™ Precast Gels (Bio-Rad). SDS-PAGE gels ran for 50 minutes at 125 V. Trans-Blot Turbo Mini 0.2 μm Nitrocellulose or polyvinylidene difluoride (PVDF) Transfer Packs and protein transfer onto membranes was achieved using the low or mixed molecular weight (MW) preprogrammed protocol in a Trans-Blot Turbo transfer system (Bio-Rad). Membranes were blocked for 1 hour at room temperature in 5% milk or 5% BSA (Millipore Sigma) in 1X Tris-buffered saline + 0.1% Tween 20 (TBST). Membranes were rinsed once with 1X TBST and primary antibodies diluted in 5% BSA in 1X TBST were added at desired concentrations. Blots were incubated in primary antibodies overnight at 4°C. The primary antibody was removed, membranes were washed 3 times in 1X TBST, and secondary antibodies diluted in 5% milk or BSA in 1X TBST were added. Blots were incubated in the secondary antibodies for 1 hour at room temperature. After incubation, membranes were rinsed 3 times with 1X TBST and the blots were incubated in Clarity ECL western blot substrate solution (Bio-Rad) for 5 minutes. Membranes were imaged using the ChemiDoc Imaging System (Bio-Rad).

### Liver tissue fixation and immunofluorescence staining

Mice were anesthetized with intraperitoneally (i.p.) injection of 250 mg/kg xylazine and 50 mg/kg ketamine. The liver was fixed by transcardial perfusion of ice-cold PBS for 2 minutes followed by ice-cold 4% paraformaldehyde (PFA) in PBS at a rate of 5 mL/min. Livers were harvested and stored in 4% PFA in PBS overnight followed by sectioning using a Leica Vibratome (50-75 μm thick). Sections were blocked with 0.5% Triton X-100 and 20% FBS in PBS for 1 hour at room temperature. Sections were incubated with the primary antibody for 1-3 days at 4°C. Tissues were washed in PBS for 1 hour and incubated with secondary antibodies overnight at 4°C. Sections were mounted with 80% glycerol in PBS.

### Histology

Paraffin-embedding, sectioning and histological staining were done by Histoserv, Inc. Histology slides were scanned and evaluated by the Molecular Pathology Unit National Cancer Institute (NCI). Three regions of interest from two mice were used for area quantification of PAS positive cells. In Image J, images were converted to 8bit, threshold was applied, and the area measured.

### Glucose and insulin tolerance tests

Fasted mice (6h) were i.p injected with 0.75 units/kg insulin (Humulin R, Lilly Pharmaceuticals) or 1 g/kg 50% Dextrose (Hospira), for insulin and glucose tolerance tests, respectively. Blood was collected from the tail vein and glucose was measured using testing strips (Ascencia Diabetes Care).

### Weight and body composition analysis

To calculate the change in body weight (BW), the final BW was subtracted from the initial BW and divided by the initial BW. Body composition (fat and lean mass) was measured in unrestrained mice by Echo MRI (EchoMRI LLC, Houston, TX) at 1:00pm.

### Food intake and energy balance analysis

Food was weighed weekly. Food intake was converted to kcal consumed/mouse/day using the energy density of the standard diet (3.1 kcal/g). Energy balance was calculated using the method described in (Ravussin et al., 2013). Briefly, the fat mass, lean mass, and body weight measurements at 4 weeks following Cre injection were subtracted from measurements taken the previous week. The difference in fat mass and lean mass values from week 3 to 4 converted to kcal (fat= 9 kcal/g, lean= 1 kcal/g). The energy gain in the body (kcal) was obtained by adding fat and lean mass values. Finally, the energy expenditure was calculated by subtracting the accumulated energy in the body from the food intake of week 4.

### Serum analysis

Kits were used according to manufacturer’s instructions to measure, FGF21 (EZRMFGF21-26K, EMD Millipore), β-hydroxybutyrate (K632, Biovision), FFA FUJIFILM (276-76491, 999-34691, 995-34791, 991-34891, 993-35191 Wako Diagnostics) and glucagon (10-1281-01, Mercodia).

### Confocal microscopy and image processing

Images were acquired using a Leica SP8 inverted confocal laser scanning microscope using 63X oil objective 1.4 NA. E-cadherin staining was used to identify PP regions of the hepatic lobule. Images were cropped contrast enhanced, background subtracted, a median filter was applied using ImageJ. Mean gray values were measured for PP and PC regions, and ratios of PC over PP were determined.

### Intravital microscopy of hepatic glucose mobilization

Mice were anesthetized with 2% isoflurane administered through a nose cone. Prior to surgery, mice were injected with heparin diluted 1:10 in 0.9% (w/v) NaCl to prevent liver ischemia. Abdominal fur was shaved, and left lobe was exposed using surgical scissors. The mouse was positioned on the stage of a Leica SP8 inverted confocal laser scanning microscope fitted with a custom stage insert. Transparent ultrasound transmission gel (Parker Aquasonic 100) was used around the incision to maintain the tissue from drying. The microscope stage was heated to 34°C and a 20X objective, 0.75 NA was used for imaging. Eighteen mg/kg of 2,000,000 MW tetramethylrhodamine-dextran (Invitrogen) was injected retro-orbitally to label the sinusoids. IVM time-lapse were acquired every 2.58 seconds for 10 minutes (512 × 512 pixels and a pixel size of 1.14 μm). After image acquisition had begun, the fluorescent glucose analog 2-(N-(7-Nitrobenz-2-oxa-1,3-diazol-4-yl)Amino)-2-Deoxyglucose (2-NBDG; 6.83 μg/kg BW, Invitrogen) dissolved in 50% dextrose (1 g/kg BW) was injected retro-orbitally. Quantification of glucose uptake and retention was performed as previously described ^24^.

### RNA isolation

Three mice per group were sacrificed and the left lobe was snap-frozen in liquid nitrogen. ∼20 mg per sample was homogenized using a TissueLyser II (Qiagen) and total RNA using the Qiagen RNeasy Mini Kit (#74104). The RNA yield was determined using a High-sensitivity RNA ScreenTape Analysis (Agilent) on an Agilent TapeStation Instrument. Samples with a RIN > 8.0 were sent for bulk mRNA sequencing.

### Bulk mRNA sequencing

TruSeq mRNA library kit (Illumina) was used to prepare the mRNA library following the manufacturer’s instructions. 200ng of total RNA was used as the input to an mRNA capture with oligo-dT coated magnetic beads. The mRNA was fragmented, and then a random-primed cDNA synthesis was performed. To prepare the library, the resulting double-strand cDNA was used as the input to a standard Illumina library prep with end-repair, adapter ligation and PCR amplification being performed. The final purified product was then quantitated by qPCR before cluster generation and sequencing. Six mRNA-Seq samples were pooled and sequenced on NextSeq using paired-end sequencing and a read length of 76 (2 × 76 cycles).

### Sequencing data analysis

Raw data files were processed using the HiSeq Real Time Analysis software (RTA 2.11.3). Files were demultiplexed and conversion of binary base calls and qualities into fastq format was performed using the Illumina bcl2fastq2.17. Cutadapt (version 1.18) was utilized to trim adapters and low-quality bases from the sequencing reads and the trimmed reads were mapped to mouse reference genome NCBI m37/mm9 and Gencode annotation M9 using STAR (version 2.7.0f) with two-pass alignment option. RSEM (version 1.3.1) was used for gene and transcript quantification.

Raw counts files were imported into the NIH Data Analysis Portal (NIDAP) for analysis in collaboration with the CCR Collaborative Bioinformatics Resource (CCBR). Genes with low counts were removed and the data was normalized by quantile normalization. Batch correction was performed on the data to account for sex differences in the samples. Differential expression of genes (DEG) analysis was performed using Limma-Voom (bioconductor-limma package v.3.38.3), and FastGSEA (bioconductor-fgsea package v.1.8.0) was used for gene set enrichment analysis (GSEA). A threshold of FDR < 0.05, FC > 1.5, FC < -1.5 was utilized to determine significance. Heatmaps were generated using hierarchical clustering on the NIDAP platform. The Venn diagrams were generated using Venny 2.1 (Oliveros, 2007). The data discussed in this publication have been deposited in NCBI’s Gene Expression Omnibus ^25^ and are accessible through GEO Series accession number GSE192404 (https://www.ncbi.nlm.nih.gov/geo/query/acc.cgi?acc=GSE192404).

### Statistical Analysis

A minimum of 3 mice were averaged per condition. As no sex by LKB1 KO interaction were observed for any of the data, male and female values were pooled. Unpaired t-tests were performed for data with equal variance and a Welch’s t-test was performed for data with unequal variance. For comparing WT and KO mice within PP or PC regions, a two-way ANOVA with Bonferroni post hoc was used, or comparing across time, a two-way repeated measures ANOVA was used. All data are shown as mean ± SEM. Statistical significance was determined by a p-value < 0.05 using GraphPad Prism (version 8.4.2). For RNA seq data, genes that had an FDR < 0.05, FC < -1.5, and FC > 1.5 were considered to be regulated by LKB1 depletion.

## Results

### LKB1 KO mice are in a prolonged fasting state resulting in a wasting phenotype

Liver-specific LKB1 KO mice were generated via tail vein injection of AAV8 Cre recombinase with a TGB promoter in LKB1 floxed mice at 6 weeks of age and analyzed 4 weeks after. Over 90% reduction in LKB1 protein was achieved (Fig 1A) with some residual LKB1 detected, likely originating from non-parenchymal cells. As previously reported in a model where LKB1 was depleted in neonates ^19^, pAMPK levels were significantly reduced in KO livers, due to loss of LKB1 which phosphorylates it (Sup Fig 1A). Overall, KO mice were smaller than WT controls which was also reflected by significant weight loss over the course of four weeks (Fig 1B and C). Weight loss occurred despite increased food intake and increase in energy expenditure, indicating poorer metabolic efficiency in KO mice (Fig 1D and E). Significant reduction in body weight from muscle and/or fat that cannot be rectified with nutritional intervention, is defined as a wasting phenotype, which is also concurrent with shifts towards catabolic hepatic gene expression and metabolism ^26^. Weight loss in KO mice included both fat and lean mass (Fig 1F and G). Additionally, substantial reduction in glycogen storage was observed in liver sections from KO mice, primarily around PC and mid-lobular regions (Fig 1H-I and Sup Fig S2). Aside for reduced glycogen storage, tissue pathology evaluation using H&E and TUNEL staining did not reveal signs of inflammation or tissue damage (Sup Fig S2). The wasting phenotype observed is consistent with previous reports in whole body ^20^ and liver-specific LKB1 KO models ^18, 19, 21^. Previously, we have shown that deletion of LKB1 at birth via Albumin Cre excision displayed a similar wasting phenotype to the one we describe here using AAV8 Cre injection in mature mice ^19^. Together, this demonstrates a vital role for liver-specific LKB1 in maintaining metabolic homeostasis in both neonate and adult mice.

**Figure 1.**
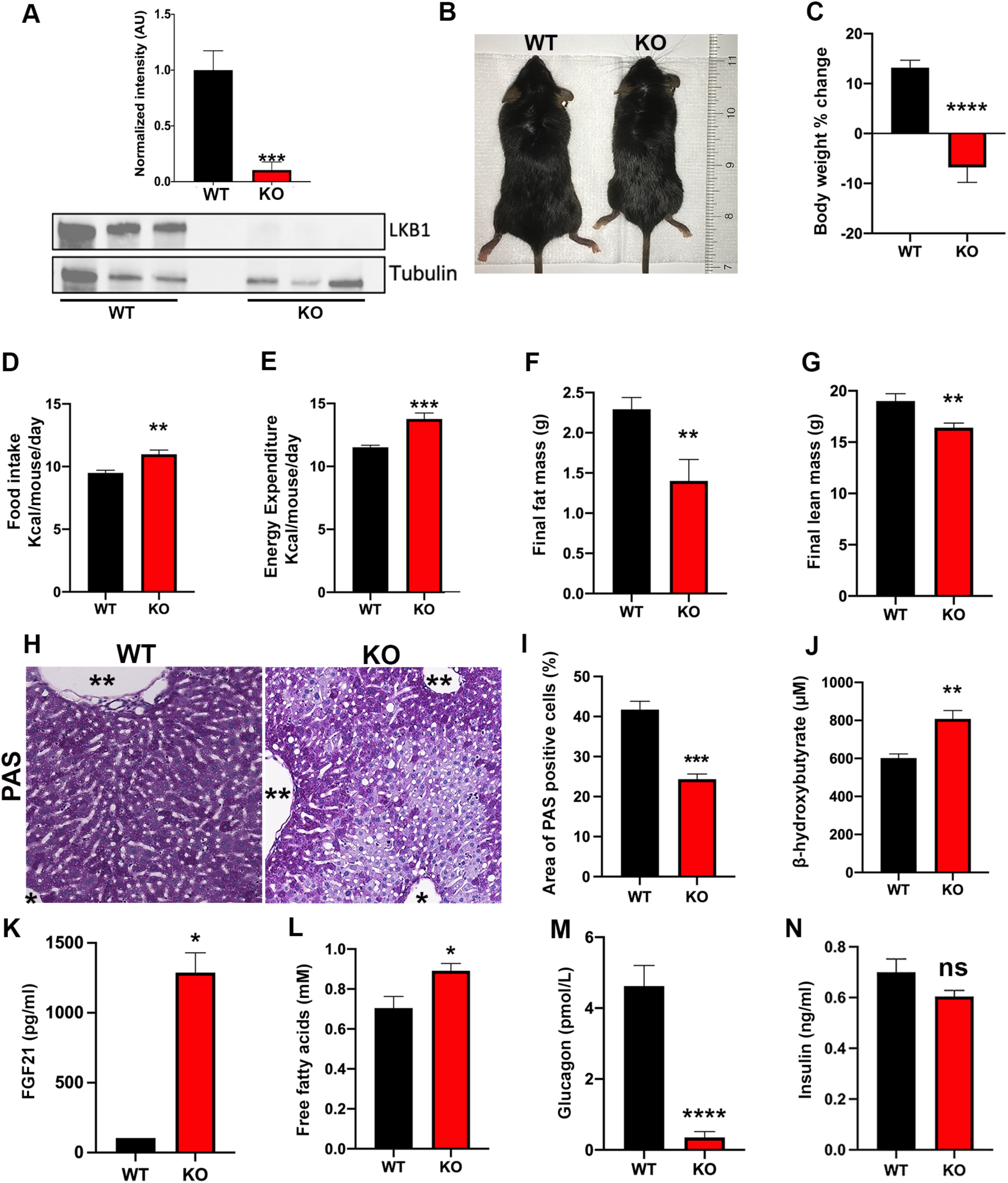
Characterization of changes in whole-body energy homeostasis in WT and KO mice. A) **Confirmation of LKB1 ablation in the liver. B)** Representative image of WT and KO mice. C) Percent change in body weight. D) Daily food intake (average values from 4 week-long measurements). E) Energy expenditure measured per mouse per day (average values from 4 week-long measurements). F) Final fat mass and G) final lean mass. H) Analysis of liver glycogen using PAS staining in liver sections from WT and KO. **Portal vessel; *Central vein. I) Quantification of PAS positive hepatocytes expressed by % area. J) Serum levels of b-hydroxybutyrate. K) Serum levels of FGF21. L) Serum levels of free fatty acids. M) Serum glucagon and N) serum insulin in WT and KO mice. In all the above experiments, N = 16-18 mice per measurement. Data are shown as mean ± SEM. P-values were calculated using an unpaired Student’s t-test. *p < 0.05, **p < 0.005, ***p < 0.0005, ****p < 0.0001, and ns= not significant.

Since LKB1 was ablated specifically in hepatocytes, we next examined whether KO livers will be a source for elevated markers of prolonged fasting/starvation. Ketone bodies are produced in the liver during extended fasting, starvation or exercise (Rui, 2014; White and Venkatesh, 2011) through fatty acid oxidation (Puchalska and Crawford, 2017). Indeed, serum level of the ketone body β-hydroxybutyrate was increased in KO mice (Fig 1J). In addition, we measured serum levels of starvation-associated hormone fibroblast growth factor 21 (FGF21) ^27^ and found it was higher in KO mice (Fig 1K). FGF21 levels have been shown to promote lipolysis in diet-induced obese mice, leading to weight loss ^28^. Additionally, FGF21 null mice are protected from muscle loss ^29^, and elevated FGF21 is associated with sarcopenia in adults ^30^. This catabolic role of FGF21 on muscle mass likely explains the reduction in muscle observed in KO mice. Consistently, serum free fatty acid levels were also elevated in KO mice (Fig 1L). Despite these elevated markers of prolonged fasting serum glucagon levels in KO mice were significantly reduced (Fig 1M), likely driven by the hyperglycemia in KOs and shown here (Fig 2A), and as previously reported ^22, 23^. On the other hand, insulin levels were unchanged (Fig 1N). This demonstrates only the liver displays a prolonged fasting phenotype, while glucagon levels apropriatly drop in response to the hyperglycemia. Collectively, these results demonstrate elevated markers indicative of hepatic prolonged fasting, consistent with the observed wasting phenotype.

**Figure 2.**
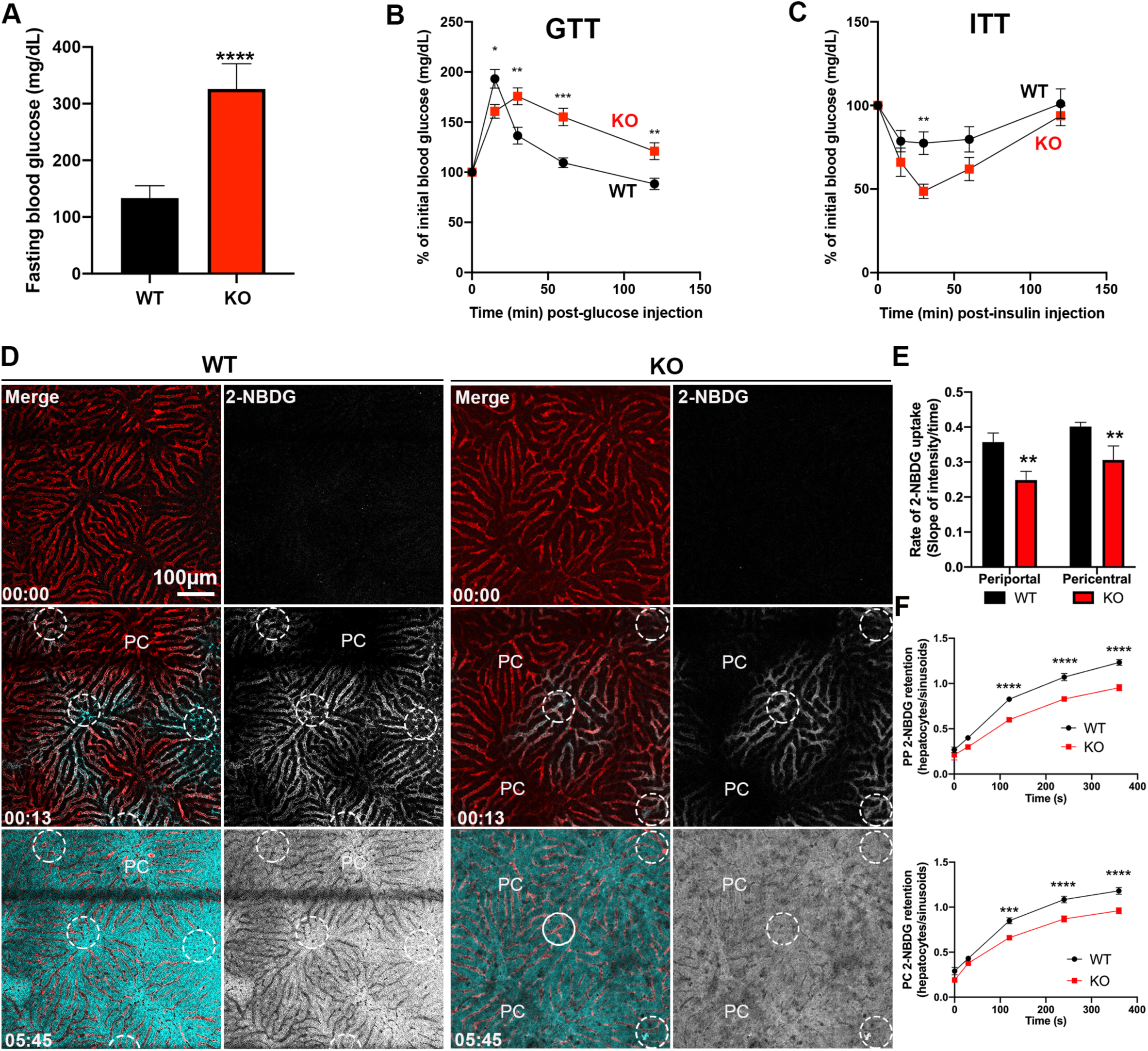
IVM of glucose uptake and retention in WT and KO mice. A) Fasting blood glucose from WT and KO mice. B) Glucose tolerance test (GTT) and C) insulin tolerance test (ITT) of WT and KO mice. D) Representative time-lapse frames of hepatic glucose uptake *in vivo* using the fluorescent glucose probe 2-NBDG (white). Mice were injected with 2-NBDG approximately 10 seconds into image acquisition. Dextran (red) was used to label the vasculature. Time = minutes: seconds. Dashed white circles indicate portal veins. CV = central vein. E) Quantification of glucose uptake rate (positive pixels/second). F) Quantification of glucose retention shown as the ratio of mean fluorescence intensity of 2-NBDG in hepatocytes to sinusoids in PP hepatocytes and PC hepatocytes. Scale bar= 100 μm. Data are shown as mean ± SEM. P-values were calculated using an unpaired Student’s t-test for fasting glucose, a two-way ANOVA for WT vs KO within PP or PC, or repeated measures two-way ANOVA for WT vs KO across time. N= 3 mice. *p < 0.05, **p < 0.005, ***p < 0.0005, and ****p < 0.00001.

### Glucose retention is reduced in LKB1 KO hepatocytes

One of the hallmark characteristics of LKB1 ablation is the dysregulation of glucose homeostasis ^22, 23^. Indeed, KO mice were severely hyperglycemic (Fig 2A) and displayed glucose intolerance (Fig 2A and B), but insulin-sensitive (Fig 2C), indicating peripheral tissues were not affected by depletion of LKB1. Given that glucose synthesis, oxidation and storage is vastly different across the liver lobule ^12^, and we observed non-uniform glycogen storage in KO mice (Fig 1G), led us to examine whether glucose uptake and retention is spatially regulated, and whether LKB1 has a role in it. To this aim, we developed an assay to measure hepatic glucose mobilization in live anesthetized mice using time-lapse IVM ^24^ using fluorescent glucose analog 2- (*N*-(7-Nitrobenz-2-oxa-1,3-diazol-4-yl)Amino)-2-Deoxyglucose (2-NBDG) (Fig 2D-F). With this approach, the rate of glucose uptake and retention in PP and PC hepatocytes was measured. In WT mice, 2-NBDG signal appeared in portal vein areas within seconds of injection and was taken up by hepatocytes over the course of 6 minutes (Fig 2D, Movie S1). Although serum glucose concentrations are reportedly higher in the portal vein ^31^, we found no difference in the rate of glucose uptake between PP and PC hepatocytes in WT mice (Fig 2E). However, in KO mice, the rate of uptake in both PP and PC hepatocytes was significantly lower compared to WT and accumulation of 2-NBDG signal was not observed in PC regions (Fig 2E). Additionally, we compared the ability of the hepatocytes to retain glucose by calculating the ratio of 2-NBDG fluorescent signal in hepatocytes over sinusoids. Hepatic glucose retention was significantly decreased in KO mice in PP and PC regions suggesting glucose is released rather than stored (Fig 2F) and consistent with the hyperglycemia and reduced glycogen. Overall, this is the first time that glucose mobilization is evaluated in a spatially resolved manner and provide novel insights to glycemic control.

### Upregulation of PP enriched genes and pathways in LKB1 KO mice

To identify the drivers of these metabolic changes in LKB1-deficient livers, we performed RNA sequencing and examined its impact on gene expression. Out of 11,669 identified genes, 1,635 were differentially expressed in the KO liver compared to WT, with roughly a 1:1 ratio of upregulated to downregulated genes (Fig 3A and Table S1). Gene set enrichment analysis (GSEA) and Ingenuity Pathway Analysis (IPA) were used to analyze the differentially expressed genes (DEG). We found 699 significantly enriched pathways, a subset of which is shown in Fig 3B. Interestingly, many pathways that are associated with PP-related functions including mitochondria oxidative phosphorylation (Sup Fig 3A and B), TCA cycle, the previously reported gluconeogenesis ^23^ and amino acids catabolism ^21^ were enriched in KO hepatocytes (Fig 3B). These pathways are typically upregulated in the fasted liver ^32^ reinforcing our observation that in the absence of LKB1 the liver is in a fasting state. Pathways associated with PC-related functions, on the other hand, such as lipogenesis and bile acid metabolism, were downregulated (Fig 3B) suggesting a bias towards PP-enriched genes in the KO liver. To investigate this idea further, we used a list of PP and PC-enriched genes compiled by Cheng *et al*. ^10^ and compared them with the DEGs identified in the LKB1 KO mice. Indeed, out of 270 PP and 461 PC genes, 89 and 157 were regulated in the LKB1 KO livers, respectively. Strikingly, within these regulated genes, there was substantial upregulation of PP-enriched and downregulation of PC-enriched genes (Fig 3C and C), suggesting the involvement of LKB1 in the spatial division of metabolic functions.

**Figure 3.**
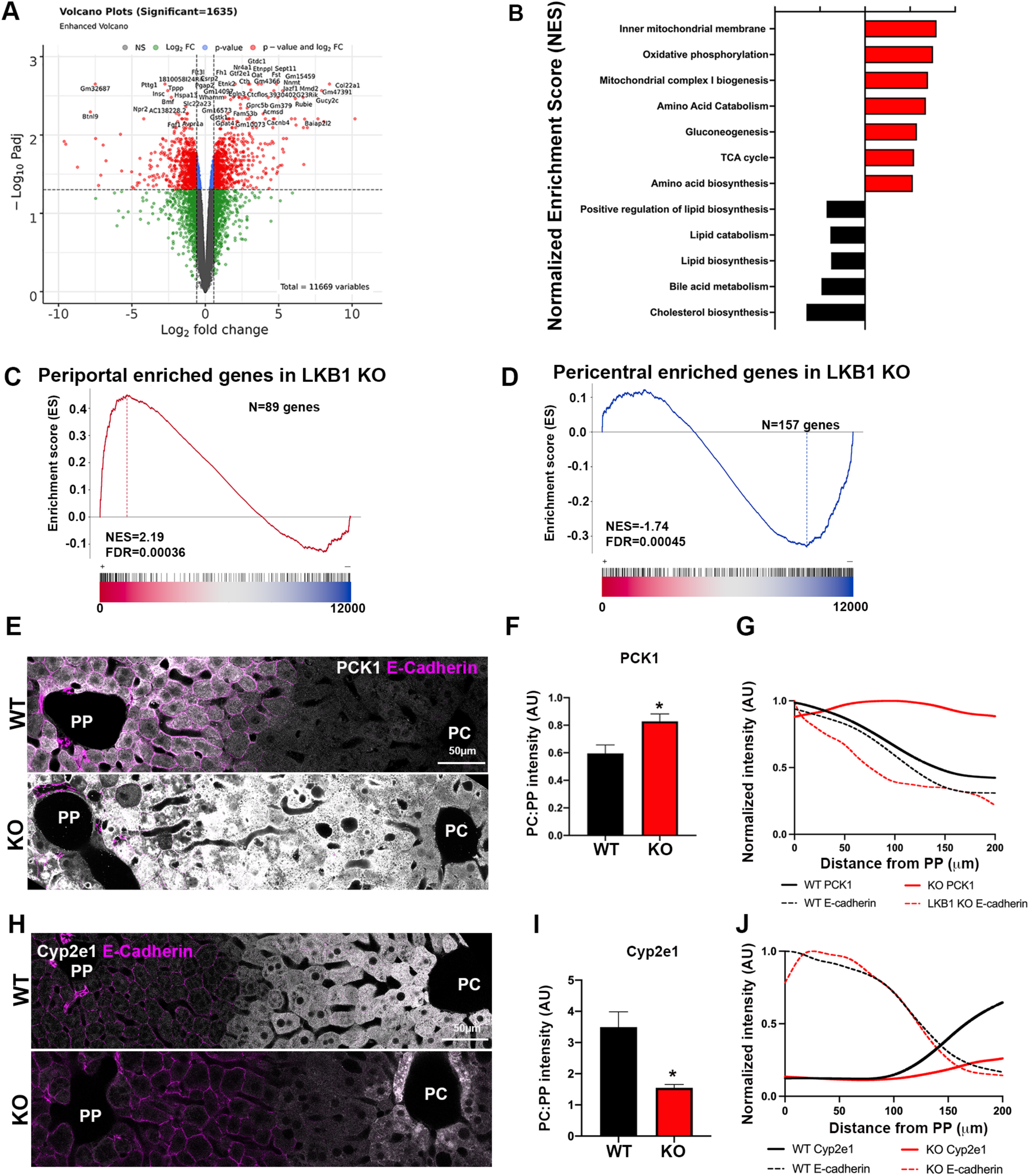
LKB1 regulates PP genes and function. A) Volcano plot showing global gene expression from WT and KO livers. N=3 mice per group. B) Enrichment of select upregulated (red) and downregulated (black) pathways in KO mice. C) Gene set enrichment plots showing expression of PP- and D) PC-associated genes, upper and lower graphs, respectively, in LKB1 KO livers. Gene list from ^10^ was used for the comparison. E) Immunofluorescence showing PCK1 (grey) localization in WT and KO liver sections. PP regions are labeled with E-cadherin (magenta). F) Quantification of PCK1 expression is shown as the ratio of PCK1 mean grey value in PC and PP hepatocytes. G) Line scan of fluorescence intensity across the lobule showing PCK1 distribution. Note that PP-enriched distribution in WT mice is lost in the absence of LKB1 and becomes uniform. H) Immunofluorescence showing Cyp2e1 (grey) localization in WT and KO liver sections. PP regions are labeled with E-cadherin (magenta). I) Quantification of Cyp2e1 expression is shown as the ratio of Cyp2e1 mean grey value in PC and PP. J) Line scan of fluorescence intensity across the lobule showing Cyp2e1 distribution. Note that PC-enriched distribution in WT mice is barely detected in the absence of LKB1. Four-8 PP or PC regions were analyzed per mouse and averaged, n=3 mice per group. Scale bar= 50 μm. Data are shown as mean ± SEM; p-values were calculated using an unpaired Student’s t-test or a Welch’s t-test.

Given these changes in zone-restricted gene expression in KO livers, immunofluorescence localization studies were performed to examine the spatial distribution of selected enzymes in liver sections. Based on gene expression, gluconeogenesis is enriched to PP hepatocytes ^12^. Consistently, protein expression of the rate-limiting enzyme Phosphoenolpyruvate Carboxykinase 1 (PCK1) was mostly confined to PP hepatocytes in WT (Fig 3E upper image). However, in the KO lobule, PCK1 expression increased and its distribution became uniform across both PP and PC hepatocytes (Fig 3E lower image). Quantification of the PC/PP fluorescence intensity ratios demonstrate the increase in PCK1 expression levels in PC regions (Fig 3F), while the line scan demonstrates the loss of PCK1 zonation in KO mice (Fig 3G, solid red line). Overall, in the WT mice PCK1 expression dramatically drops and therefore barely detected in PC regions, whereas in the KO mice it is present across the lobule suggesting a loss of functional zonation (Fig 1G). Similar results were obtained with the PP urea cycle enzyme Arginase 1, (Sup Fig 3C-E). The spatial localization of PC enzymes was also investigated, including Cyp2e1 which is involved in xenobiotic metabolism. Cyp2e1 is mostly confined to PC hepatocytes in the WT, and this pattern was lost in KO due to reduced expression levels of the enzyme and restricted distribution around the central vein (Fig 3H-J). Similar results were obtained with the PC lipogenic enzyme acetyl-CoA carboxylase 1 (ACC1), (Sup Fig 3F-H). Overall, in the absence of LKB1, the liver defaults to a PP metabolic program that overrides PC-related gene expression and functions. This loss in zonation and new uniform distribution of PP enzymes indicates that LKB1 signaling is important in maintaining liver zonation.

### LKB1 regulates glucagon target genes, but not Wnt/b-catenin

To understand how LKB1 affects liver zonation IPA analysis was performed to identify potential regulators based on the DEG. Interestingly, glucagon was identified as the top upstream activator of LKB1 (Fig 4A). Since glucagon was reported to regulate liver zonation by promoting the expression of PP genes ^10^ we next investigated the relationship between LKB1- and glucagon-regulated genes. To this aim, DEG identified in LKB1 KO was compared with those identified in glucagon KO mice ^10^. Glucagon KO mice were selected for this comparison because, like the LKB1 KO, they displayed low serum glucagon. Of the 297 DEGs identified in glucagon KO livers, 40% overlapped with DEGs in LKB1 KO (Fig 4B and Table S2). Interestingly, the overlapping genes inversely correlated (Fig 4C) suggesting that LKB1 has an overall inhibitory impact on glucagon-related gene expression. In glucagon KO mice, PC genes were upregulated ^10^. In LKB1 KO mice, despite reduced glucagon levels (Fig 1M), PP-enriched genes were enhanced, thus reinforcing the notion that LKB1 negatively regulates glucagon-related transcriptional changes. Similar analysis of Wnt target genes revealed that out of the 1003 DEG, 20% of the LKB1 KO overlapped (Fig 4D and Table S3). However, there was no correlation in expression between the overlapping genes (Fig 4E). These results suggest that upregulation in PP-enriched and decreased PC-enriched gene expression in LKB1 deficient hepatocytes is driven by uninhibited glucagon-related pathways rather than changes to Wnt/b-catenin. To confirm this, we also compared DEGs from LKB1 KO with those reported in liver-specific glucagon receptor KO mice ^33^. Out of the 1744 DEGs identified in glucagon receptor KO livers, 15% overlapped with DEGs in LKB1 KO (Sup Fig S4 and Table S4) and their expression displayed a significant inverse correlation (Fig S4A and B). The common affected genes in LKB1 KO, glucagon KO and glucagon receptor KO are shown in Fig 4F and Table S5. We posit that LKB1 acts as a brake for glucagon-related gene expression. Once that brake is gone (i.e. LKB1 KO), the negative feedback loop is broken, allowing glucagon-related pathways to increase unregulated. This notion is reinforced by the fact that glucagon-related genes are elevated in the face of low serum glucagon levels in LKB1 KO mice.

**Figure 4.**
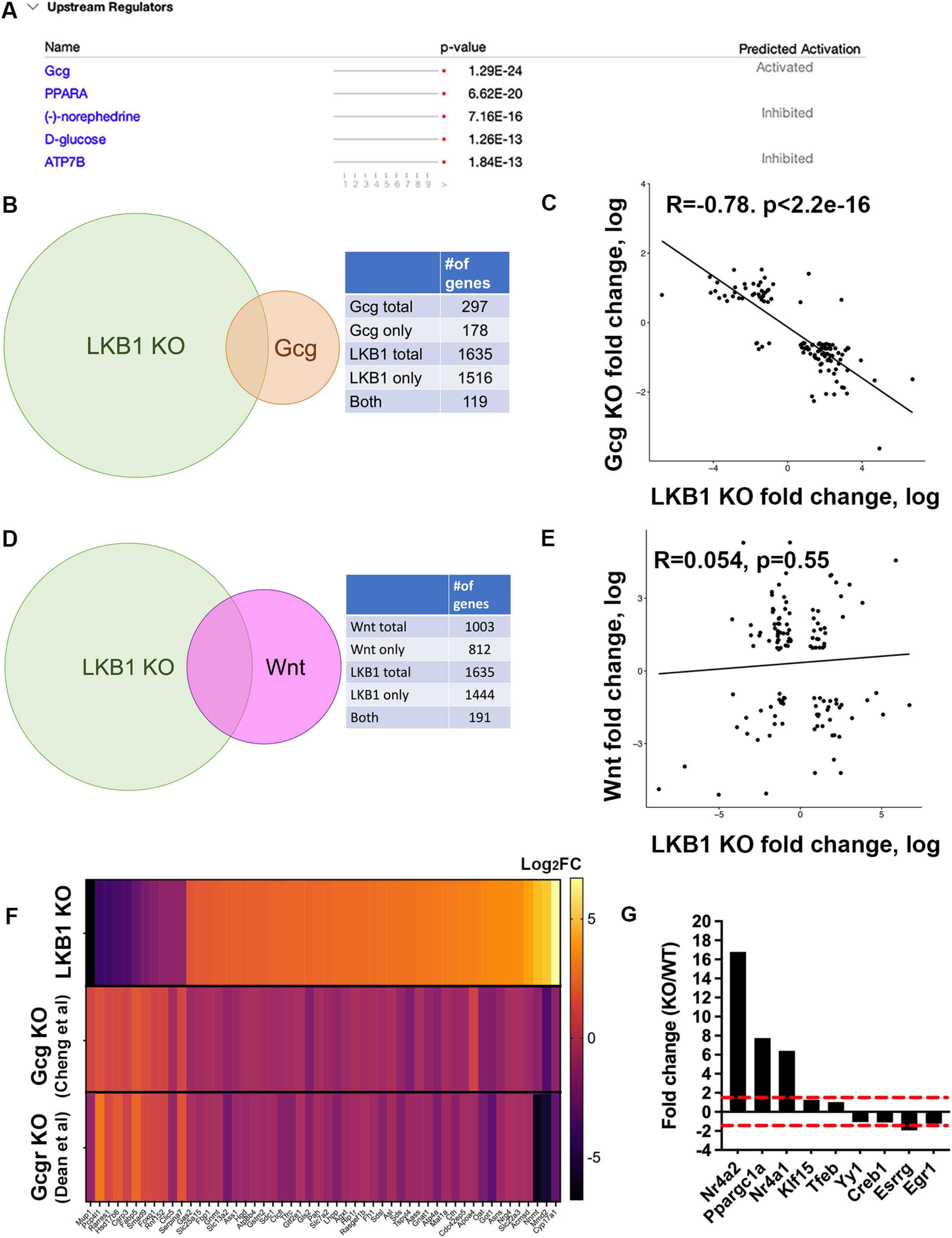
LKB1 negatively regulates glucagon-induced genes, but not Wnt-regulated genes. A) Upstream regulators of LKB1 identified using Ingenuity Pathway Analysis (IPA). B) Venn diagram of differentially expressed genes in LKB1 KO (current study) and glucagon KO ^10^ showing 119 overlapping genes. C) The correlation between LKB1 KO and glucagon KO genes. D) Venn diagram of differentially expressed genes in LKB1 KO (current study), and Wnt-regulated ^42, 50^ showing 191 overlapping genes. E) The correlation between LKB1 KO and Wnt-regulated genes. F) Heat map of representative differentially expressed genes in LKB1 KO, Gcg KO ^10^and Gcgr KO ^33^ G) Fold change of transcription factors in WT versus LKB1 KO mice.

Glucagon is involved in the hepatic fasting response through activation of transcription factors. Initially, glucagon stimulates cAMP/CREB pathway which leads to expression of additional transcription factors and a second wave of gene expression facilitating the temporal regulation of the fasting response ^8^. Therefore, expression levels of transcription factors involved in the hepatic fasting response in the LKB1 KO were examined. Significant increases in Nr4a1 (6-fold) and Nr4a2 (16-fold) were found in KO mice (Fig 4G). These factors are involved in upregulating gluconeogenesis and the expression of FGF21 ^34, 35^. Additionally, increased expression of Ppargc1a (7-fold) was observed which was previously described in LKB1 KO mice ^22^, and is involved in FAO and ketogenesis ^36^. These transcription factors are part of the delayed response to glucagon and may explain the exacerbated and prolonged hepatic fasting phenotype (Fig 1). Lastly, Figure 5 summarizes our findings in a model. We propose a negative feedback loop in which LKB1 keeps the glucagon-related hepatic fasting response under control. Depletion of LKB1 results in unrestrained activation of glucagon-related pathways including gluconeogenesis, glucose efflux, ketogenesis and lipolysis. These zone-restricted functions extend across the hepatic lobule in mutant mice leading to loss of liver zonation. These findings explain the wasting phenotypes resulting in overall weight loss, inability to store fuels and metabolic inefficiency. Additionally, this study highlights the possibility that glucagon-related hepatic fasting is not only transcriptionally controlled in time ^8^, but also spatially regulated by LKB1 across the hepatic lobule.

**Figure 5.**
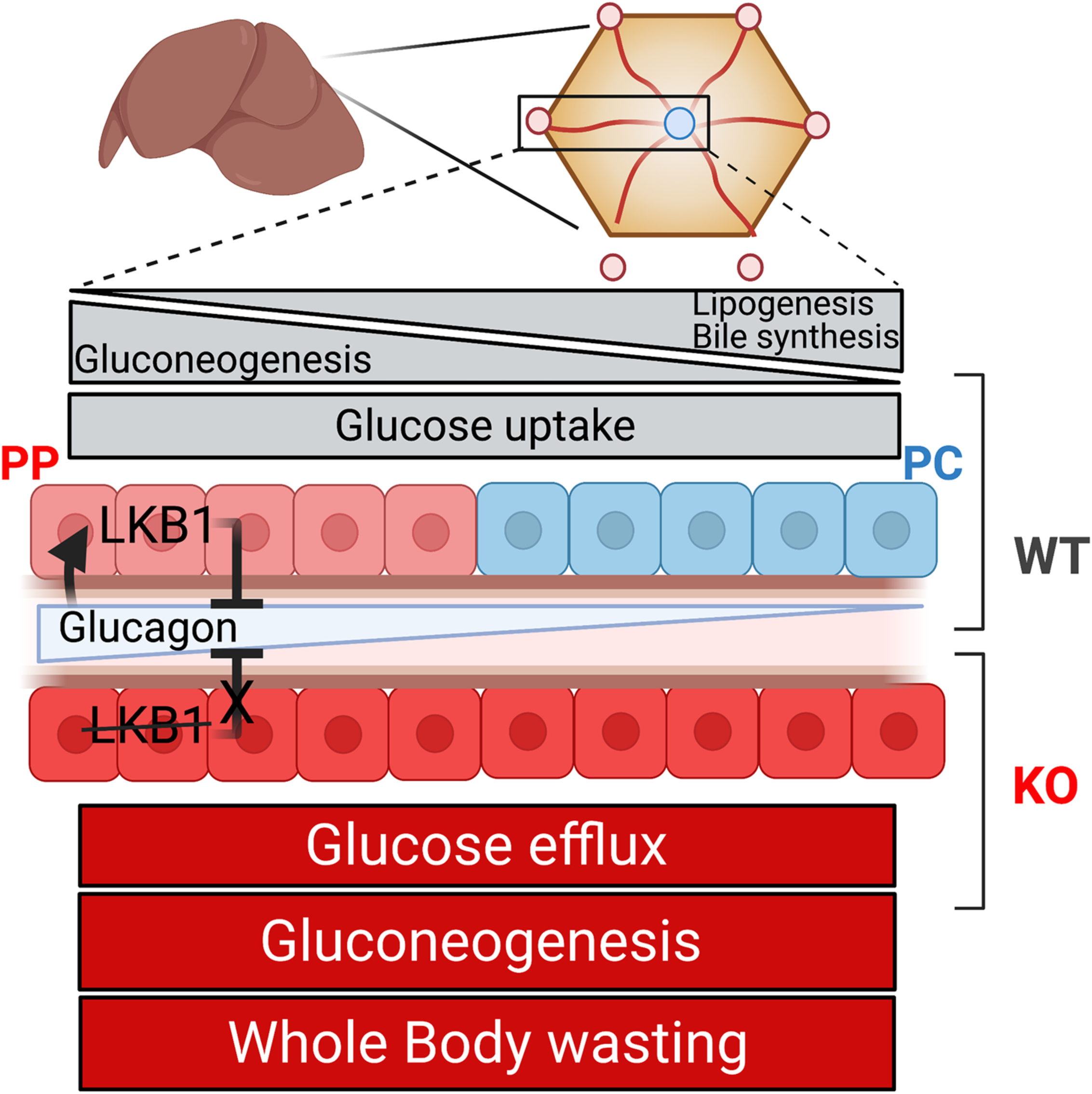
Graphical summary of results, showing how loss of LKB1 allows glucagon genes to be up regulated leading to metabolic inefficiency and periportalization of the lobule.

## Discussion

Fasting is an essential adaptive response to simultaneously mobilize nutrients as a fuel source while conserving energy expenditure. During diseases such as NAFLD and type 2 diabetes, an inability to shut down this fasting response results in persistent gluconeogenesis and lipolysis ^37^, despite adequate nutrient consumption. This excessive mobilization of fuels worsens the inability to arrest the fasting response, and further exacerbates disease progression. Here, we show that LKB1 is vital for the inhibition of the hepatic fasting response. Our findings that LKB1 KO mice have a wasting phenotype (Fig 1) despite increased food intake (Fig 1D) and circulating substrates in the blood (Fig 1J-N and 2A) are indicative of disruption in the homeostatic control of energy balance and suggestive of overall metabolic inefficiency. Energy expenditure allows the evaluation of overall metabolic efficiency by measuring the ability to convert ingested energy into fat and protein ^38^. In an attempt to maintain bodyweight during fasting, energy expenditure is decreased by reducing physical activity ^39^ and diet-induced thermogenesis ^40^. Supporting this, Just and colleagues previously showed that liver specific LKB1 KO had decreased locomotor activity ^21^. As expected, we measured significantly higher energy expenditure in KO mice despite being in a phenotypical hepatic fasting state (Fig 1E), reflecting metabolic inefficiency in LKB1 KO mice. Although we do not know the cause for this inefficiency, the dramatic reduction in hepatocyte heterogeneity reported here may lead to impaired division of liver function, causing reduced efficiency. Alternatively, it is possible that due to the severe hyperglycemia in KO mice, the diet-induced thermogenesis is increased.

One of the key phenotypes described in LKB1 KO mice is dysregulated glucose homeostasis ^21-23^. Hepatocytes shift between glucose uptake for storage and efflux during fed and fasted states, respectively. Despite the fact that glucose uptake, storage and efflux are differentially regulated across the hepatic lobule ^12, 13^, it is not clear if LKB1 is involved and how. To measure the changes in glucose uptake and release, we developed an IVM-based assay to measure glucose mobilization in space and time ^24^. We measured markedly reduced glucose uptake and retention in KO hepatocytes indicating they are in constitutive efflux state (Fig 2D-F). These results are consistent with the severe hyperglycemia and glucose intolerance (Fig 2A and B), reduced glycogen storage (Fig 1H and I), and upregulation of gluconeogenesis genes (Fig 3B-E). Interestingly, Berndt and colleagues used modeling to propose that PP hepatocytes release and PC hepatocytes take up glucose ^41^. However, *in vivo*, we did not detect any difference in the rate of glucose uptake and/or retention between PP and PC hepatocytes in WT or KO (Fig 2D and E). Similar kinetics in PP and PC are likely achieved through differential expression of high- and low-affinity glucose transporters across the lobule ^12^ and redistribution of these transporters when glucose availability shifts. This is the first time that LKB1 has been shown to play a vital role in the spatial metabolism of glucose in the liver. Our results underscore the importance of measurements of physiological parameters in the intact tissue.

This unique spatial organization of hepatocytes (Fig 5 and Sup Fig 1E) results in the formation of nutrient, oxygen, hormone, and morphogen gradients, which in turn, shape hepatocytes’ gene expression. Indeed, a recent study employing single-cell RNA sequencing demonstrated that 50% of liver genes are zonated ^42^. Key regulators of this zonation were identified and include Wnt/b-catenin, glucagon, hypoxia, H-ras, and growth and thyroid hormones ^12^. However, these do not account for all the zonated genes, highlighting the existence of additional regulators of zonation. Our findings show that LKB1 plays a significant role in liver zonation. In the absence of LKB1, the spatial division of hepatic metabolism is abolished. Instead, we observed a striking upregulation of PP-associated genes, expansion of their spatial distribution and downregulation of PC-related genes (Fig 3B-D and Sup Fig 3A and B). In a previous study, Cheng *et al*. showed that glucagon oppose Wnt/b -catenin activity by inducing PP gene expression ^10^. Our analysis identified glucagon as a probable upstream activator of LKB1. Furthermore, we show a strong inverse correlation between glucagon and LKB1 regulated genes. This apparent interplay between glucagon and LKB1 indicates a possible fine tuning of the hepatocyte’s response to glucagon, although additional studies are warranted to confirm this. We propose a negative feedback loop between glucagon and LKB1, leading to fine tuning of hepatocyte’s response to glucagon. Indeed, Cheng et al. showed that glucagon ablation led to downregulation of PP associated pathways and upregulation of PC ones, reinforcing the notion that LKB1 influences glucagon signaling ^43^. This type of regulation is reminiscent of the balancing act between Wnt/b-catenin and its negative regulator Adenomatous Polyposis Coli Protein (APC) in the regulation of PC genes ^44, 45^. Previous reports also indicated a cross talk between LKB1 and Wnt signaling in different models ^46-48^. However, our data shows no correlation between the expression of Wnt targets and LKB1 regulated genes (Fig 4D and E) suggesting that the effects of LKB1 on zonation are largely independent of Wnt. While mechanistic studies to elucidate how LKB1 counterbalances glucagon are warranted, we propose that hepatocyte nutrient sensing through LKB1, together with extrinsic factors such as glucagon, orchestrate the patterned gene expression observed in the hepatic lobule. This study is focused on glucagon, however, we cannot rule out other factors such as corticosterone and epinephrin could contribute to the phenotype.

Our study demonstrates that loss of LKB1 in the liver leads to a prolonged hepatic fasting response, which contribute to weight loss, decreased glycogen, lean mass, and adipose tissue reserve and an overall decrease in metabolic efficiency (Fig 1). For the first time, we identify LKB1 as a vital inhibitor of the glucagon-related hepatic fasting response to prevent a wasting phenotype. We posit that LKB1 acts as an important brake in the negative feedback loop of glucagon signaling in the liver. One possibility is that LKB1 interact with transcription factors in the nucleus and affects their activity. It was previously shown that the transcription factor Nr4a1 (Nur77) binds and sequesters LKB1 in the nucleus, thereby attenuating AMPK activation ^49^. In turn, this interaction may also regulate the activity of transcription factors and, like shown in this study, lead to an overall upregulation of glucagon-related genes. Attenuation of the glucagon response via metformin and GLP-1 agonists has become one of the most effective drug targets for obesity, type 2 diabetes, and NAFLD ^4, 6^. Therefore, increasing our understanding the regulatory mechanisms behind glucagon signaling is vital to improve new pharmacological therapeutics. Here, the wasting phenotype caused by uncontrolled glucagon-related gene expression in the liver was observed alongside complete ablation of metabolic zonation in the hepatic lobule, raising the provocative possibility that the fasting response is not only regulated transcriptionally in time ^8^ but also in space within the hepatic lobule. This could be critical for the organism’s survival in sustaining the fasting response if nutrients are unavailable. The lobule’s anatomy allows for increased capacity of PP-like hepatocytes and glucagon-LKB1 signaling axis may be at the nexus of the spatio-temporal regulation of the fasting response. While confirming previous reports of body weight loss (^19-21^) and reduced muscle mass ^21^ in LKB1 KO mice, this paper extends these findings to demonstrate a role for LKB1 in modulating the hepatic fasting response through glucagon-induced genes and overall impact on liver zonation, Further, we show for the first time that the disruption in glucose homeostasis in LKB1 KO mice with spatial and temporal resolution. Given the conserved role of LKB1 in cell polarity, it is possible it is also involved in the organization of the hepatic lobule which represents polarity in the tissue scale. Mechanistically, it is unclear how these zone-specific changes influence metabolic efficiency seen in LKB1 KO mice. Future studies that attempt to restore liver zonation and rescue the wasting phenotype in LKB1 KO mice would identify its disruption as the primary cause of metabolic inefficiency in these mice. Nonetheless, this study reveals a new mechanism by which hepatic metabolic compartmentalization is regulated by nutrient sensing.

## Contact information

Natalie Porat-Shliom, Center for Cancer Research, National Cancer Institute. Building 10, Room 12C207 MSC 1919, Bethesda, MD 20892. Phone: 240-760-6847.

## Abbreviations

LKB1: Liver Kinase B1
NAFLD: Non-Alcoholic Fatty Liver Disease
GLP-1: Glucagon-like peptide 1
PP: Periportal
PC: Pericentral
AMPK: AMP-activated Protein Kinase
ARK: AMPK-related
KO: Knock out
WT: Wild Type
PCK1: phosphoenolpyruvate carboxykinase 1
GAPDH: Glyceraldehyde 3-phosphate dehydrogenase
AAV8: Adeno-Associated Virus 8
TBS: Tris-buffered saline
TBST: Tris-buffered saline triton
PBS: Phosphate-buffered saline
FBS: Fetal Bovine serum
PFA: Paraformaldehyde
I.p.: intraperitoneally
BW: Body weight
FGF21: Fibroblast growth factor 21
ACC1: Acetyl-CoA carboxylase
NaCl: Sodium Chloride
2-NBDG: 2-(N-(7-Nitrobenz-2-oxa-1,3-diazol-4-yl)Amino)-2-Deoxyglucose
PCR: Polymerase chain reaction
qPCR: Quantitative polymerase chain reaction
DEG: Differential expression of genes
GSEA: Gene set enrichment analysis
NIDAP: NIH Data Analysis Portal
IPA: Ingenuity pathway analysis
FC: Fold change
FDR: False discovery rate

## Financial Support

This work was supported by the Intramural Research Program at the NIH, National Cancer Institute (1ZIABC011828). The authors have no conflicts to report.

## Acknowledgment

We thank members of the Porat-Shliom lab, Drs. Will Printz, Win Arias and Julie Donaldson for critical reading of the manuscript. Dr. Oksana Gavrilova from the NIDDK Mouse Metabolism Core for mouse phenotyping studies. CCR Sequencing Facility at the Frederick National Laboratory for Cancer Research and Dr. Alexei Lobanov from the The CCR Collaborative Bioinformatics Resource (CCBR).

## Supplementary Figures

**Figure S1.**
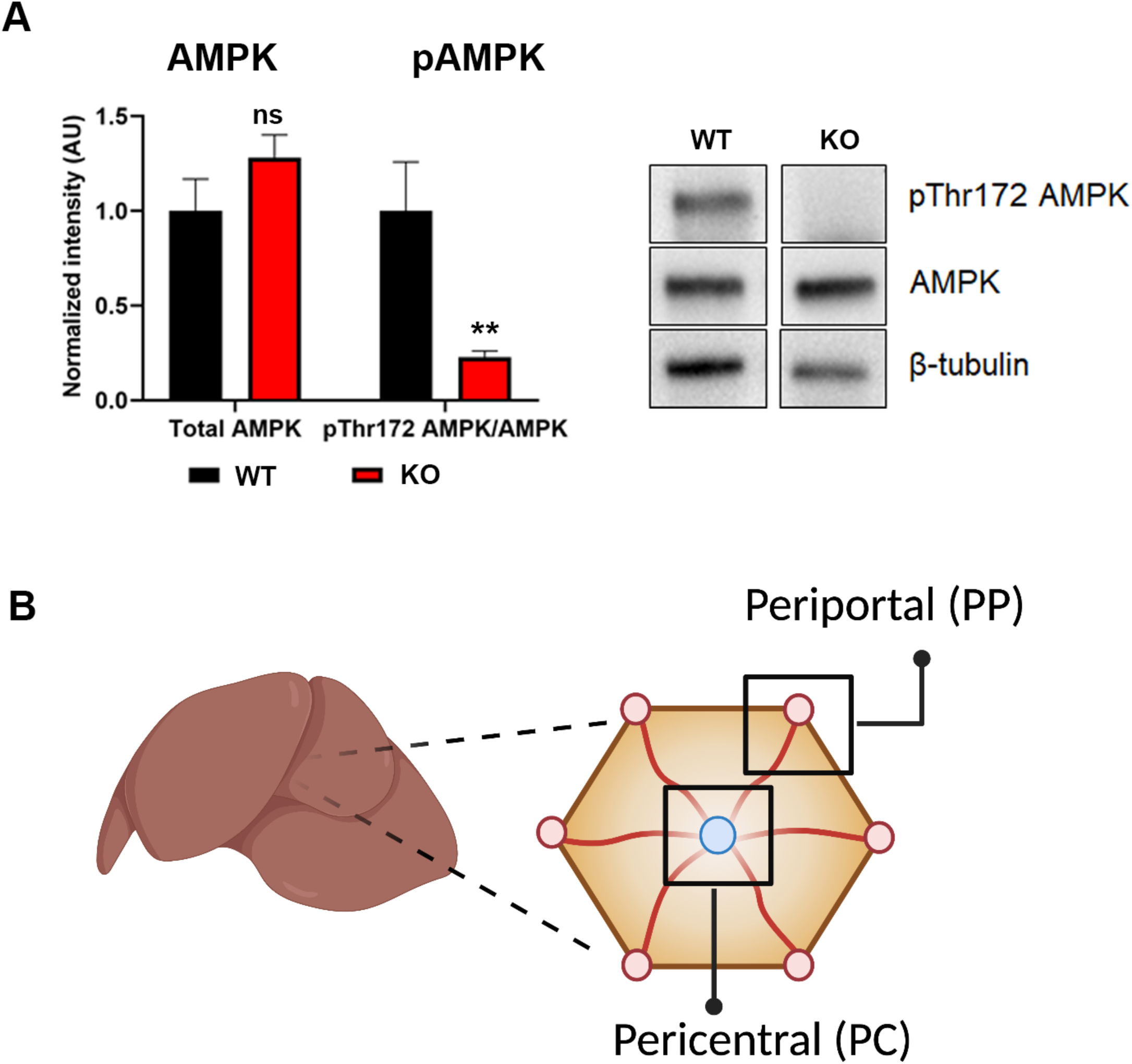
Confirmation of reduce pAMPK levels in LKB1 KO. A) Quantification of pAMPK leves in WT and KO mice and a representative western blot showing total AMPK, pAMPK (Thr172) and b-tubulin. B) Schematic visualization of the hepatic lobule. Data are shown as mean ± SEM; p-values were calculated using an unpaired Student’s t-test or a Welch’s t-test. *p < 0.05, **p < 0.005, ***p < 0.0005, and ****p < 0.00001.

**Figure S2.**
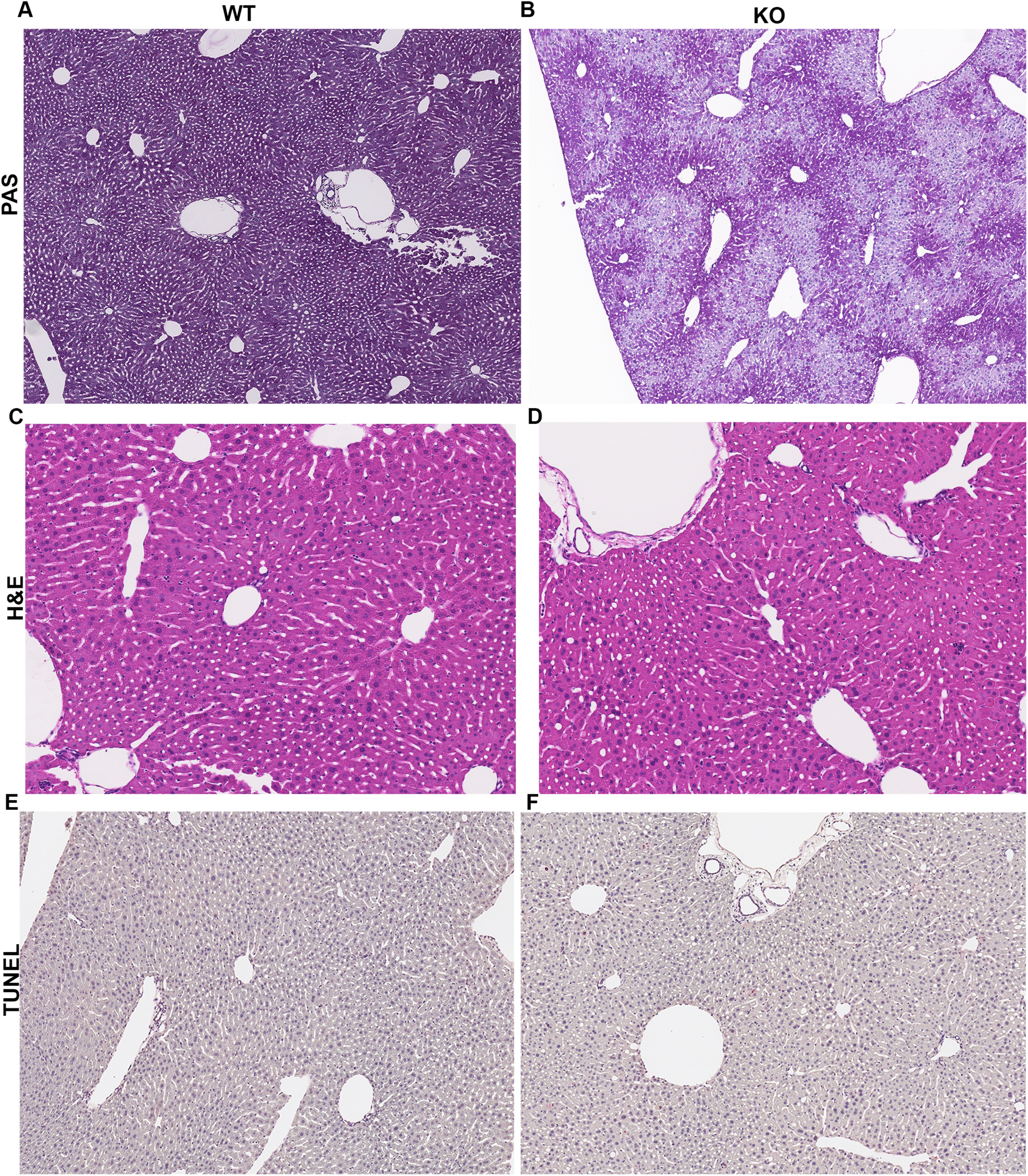
Impaired glycogen storage in LKB1 KO mice. A) Liver glycogen as demonstrated by PAS staining (purple-red, 100X). PAS-positive cells were evenly distributed in WT mice, but were significantly reduced in KO mice, particularly in PC areas. B) H&E and C) TUNEL staining show otherwise similar histology and no indication of inflammation or injury between WT and KO mice. Magnification 100X.

**Figure S3.**
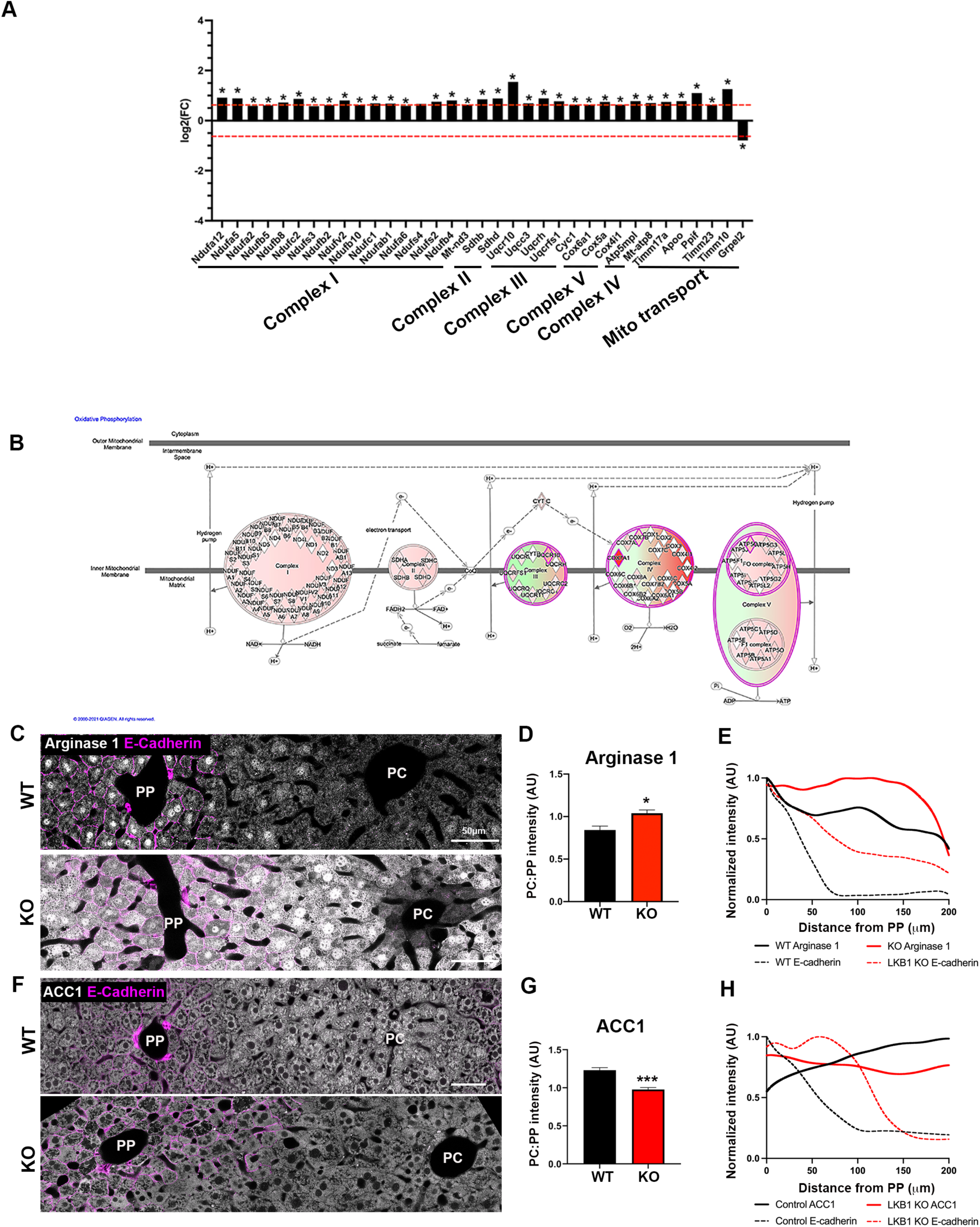
Up regulation of PP genes and function in LKB1 KO. A) Fold change of genes involved in oxidative phosphorylation and mitochondrial transport in WT and KO mice. B) IPA representation of changes in oxidative phosphorylation genes in KO mice. C) Immunofluorescence showing Arginase 1 (grey) localization in WT and KO liver sections. PP regions are labeled with E-cadherin (magenta). D) Quantification of Arginase 1 expression is shown as the ratio between mean grey value in PC and PP hepatocytes. E) Line scan of fluorescence intensity across the lobule showing Arginase 1 distribution. Note that PP-enriched distribution in WT mice is lost in the absence of LKB1 and becomes uniform. H) Immunofluorescence showing ACC1 (grey) localization in WT and KO liver sections. PP regions are labeled with E-cadherin (magenta). I) Quantification of ACC1 expression is shown as the ratio between mean grey value in PC and PP hepatocytes. J) Line scan of fluorescence intensity across the lobule showing ACC1 distribution. Note that PC-enriched distribution in WT mice is lost in the absence of LKB1 and becomes uniform. 4-8 veins were analyzed per mouse and averaged, n=3 mice per group. Scale bar= 50 μm. All data are shown as mean ± SEM, n= 3-5 mice. P-values were calculated using an unpaired Student’s t-test (Prism 8.4.2), *p-value<0.05, **p-value=0.005, ***p-value=0.0005.

**Figure S4.**
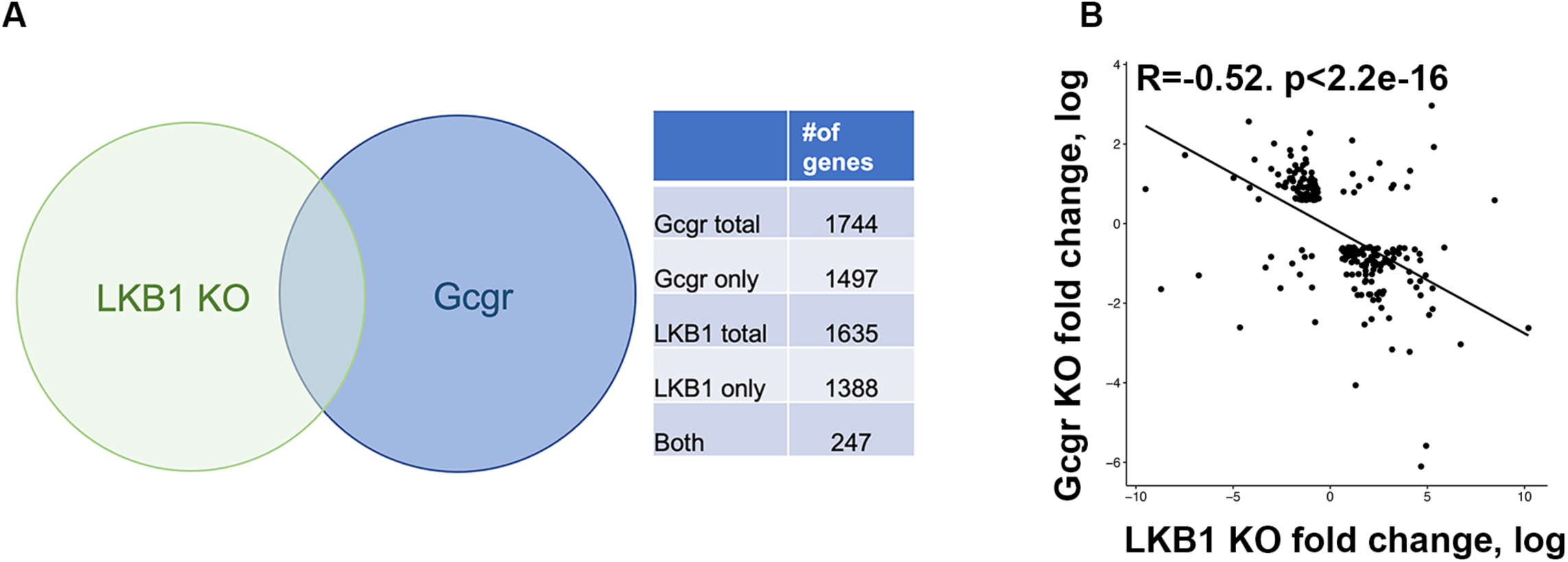
Comparison between differentially expressed genes identified in LKB1 KO and glucagon receptor KO. A) Venn diagram of differentially expressed genes in LKB1 KO (current study) and liver specific glucagon receptor KO ^33^ showing 247 overlapping genes. B) The correlation between overlapping genes from LKB1 KO and glucagon receptor KO.

## Supplementary tables

Table S1_LKB1 KO DEGs

Table S2_LKB1 KO and Gcg KO comparison

Table S3_LKB1 KO and APC KO comparison

Table S4_LKB1 KO and Gcgr KO comparison

Table S5_LKB1 KO, Gcg KO and Gcgr KO comparison

## Supplementary Movies

Movie S1. **IVM of 2-NBDG uptake in WT liver**. Vasculature was labeled by dextran (red) which was injected prior to imaging and 2-NBDG is shown in green, which was injected after the first image was acquired. Images were acquired every 2.5 seconds for 5 minutes.

Movie S2. **IVM of 2-NBDG uptake in KO liver**. Vasculature was labeled by dextran (red) which was injected prior to imaging and 2-NBDG is shown in green, which was injected after the first image was acquired. Images were acquired every 2.5 seconds for 5 minutes.

## Notes

https://www.ncbi.nlm.nih.gov/geo/query/acc.cgi?acc=GSE192404

### Competing Interest Statement

The authors have declared no competing interest.

### Summary of Updates

Figures 1,3 and 4 revised. Sup Fig 1,2,3 and 4 revised. Additional gene comparisons were performed.

## References

1. Jiang, G.; Zhang, B. B., Glucagon and regulation of glucose metabolism. Am J Physiol Endocrinol Metab 2003, 284 (4), E671–8.

2. Samuel, V. T.; Shulman, G. I., Nonalcoholic Fatty Liver Disease as a Nexus of Metabolic and Hepatic Diseases. Cell Metab 2018, 27 (1), 22–41.

3. Seppala-Lindroos, A.; Vehkavaara, S.; Hakkinen, A. M.; Goto, T.; Westerbacka, J.; Sovijarvi, A.; Halavaara, J.; Yki-Jarvinen, H., Fat accumulation in the liver is associated with defects in insulin suppression of glucose production and serum free fatty acids independent of obesity in normal men. J Clin Endocr Metab 2002, 87 (7), 3023–3028.

4. Rosenstock, J.; Wysham, C.; Frias, J. P.; Kaneko, S.; Lee, C. J.; Fernandez Lando, L.; Mao, H.; Cui, X.; Karanikas, C. A.; Thieu, V. T., Efficacy and safety of a novel dual GIP and GLP-1 receptor agonist tirzepatide in patients with type 2 diabetes (SURPASS-1): a double-blind, randomised, phase 3 trial. Lancet 2021, 398 (10295), 143–155.

5. Drucker, D. J., Mechanisms of Action and Therapeutic Application of Glucagon-like Peptide-1. Cell Metab 2018, 27 (4), 740–756.

6. Wilding, J. P. H.; Batterham, R. L.; Calanna, S.; Davies, M.; Van Gaal, L. F.; Lingvay, I.; McGowan, B. M.; Rosenstock, J.; Tran, M. T. D.; Wadden, T. A.; Wharton, S.; Yokote, K.; Zeuthen, N.; Kushner, R. F.; Group, S. S., Once-Weekly Semaglutide in Adults with Overweight or Obesity. N Engl J Med 2021, 384 (11), 989.

7. Miller, R. A.; Chu, Q. W.; Xie, J. X.; Foretz, M.; Viollet, B.; Birnbaum, M. J., Biguanides suppress hepatic glucagon signalling by decreasing production of cyclic AMP. Nature 2013, 494 (7436), 256–260.

8. Goldstein, I.; Hager, G. L., The Three Ds of Transcription Activation by Glucagon: Direct, Delayed, and Dynamic. Endocrinology 2018, 159 (1), 206–216.

9. Ravnskjaer, K.; Madiraju, A.; Montminy, M., Role of the cAMP Pathway in Glucose and Lipid Metabolism. Handb Exp Pharmacol 2016, 233, 29–49.

10. Cheng, X.; Kim, S. Y.; Okamoto, H.; Xin, Y.; Yancopoulos, G. D.; Murphy, A. J.; Gromada, J., Glucagon contributes to liver zonation. Proc Natl Acad Sci U S A 2018, 115 (17), E4111–E4119.

11. Jungermann, K.; Sasse, D., Heterogeneity of Liver Parenchymal-Cells. Trends Biochem Sci 1978, 3 (9), 198–202.

12. Ben-Moshe, S.; Itzkovitz, S., Spatial heterogeneity in the mammalian liver. Nat Rev Gastroenterol Hepatol 2019, 16 (7), 395–410.

13. Cunningham, R. P.; Porat-Shliom, N., Liver Zonation -Revisiting Old Questions With New Technologies. Front Physiol 2021, 12, 732929.

14. Kietzmann, T., Metabolic zonation of the liver: The oxygen gradient revisited. Redox Biology 2017, 11, 622–630.

15. Preziosi, M.; Okabe, H.; Poddar, M.; Singh, S.; Monga, S. P., Endothelial Wnts regulate beta-catenin signaling in murine liver zonation and regeneration: A sequel to the Wnt-Wnt situation. Hepatol Commun 2018, 2 (7), 845–860.

16. Fu, D.; Wakabayashi, Y.; Ido, Y.; Lippincott-Schwartz, J.; Arias, I. M., Regulation of bile canalicular network formation and maintenance by AMP-activated protein kinase and LKB1. Journal of Cell Science 2010, 123 (19), 3294–3302.

17. Fu, D.; Wakabayashi, Y.; Lippincott-Schwartz, J.; Arias, I. M., Bile acid stimulates hepatocyte polarization through a cAMP-Epac-MEK-LKB1-AMPK pathway. P Natl Acad Sci USA 2011, 108 (4), 1403–1408.

18. Homolya, L.; Fu, D.; Sengupta, P.; Jarnik, M.; Gillet, J. P.; Vitale-Cross, L.; Gutkind, J. S.; Lippincott-Schwartz, J.; Arias, I. M., LKB1/AMPK and PKA control ABCB11 trafficking and polarization in hepatocytes. PLoS One 2014, 9 (3), e91921.

19. Porat-Shliom, N.; Tietgens, A. J.; Van Itallie, C. M.; Vitale-Cross, L.; Jarnik, M.; Harding, O. J.; Anderson, J. M.; Gutkind, J. S.; Weigert, R.; Arias, I. M., Liver Kinase B1 Regulates Hepatocellular Tight Junction Distribution and Function In Vivo. Hepatology 2016, 64 (4), 1317–1329.

20. Shan, T.; Xiong, Y.; Kuang, S., Deletion of Lkb1 in adult mice results in body weight reduction and lethality. Sci Rep 2016, 6, 36561.

21. Just, P. A.; Charawi, S.; Denis, R. G. P.; Savall, M.; Traore, M.; Foretz, M.; Bastu, S.; Magassa, S.; Senni, N.; Sohier, P.; Wursmer, M.; Vasseur-Cognet, M.; Schmitt, A.; Le Gall, M.; Leduc, M.; Guillonneau, F.; De Bandt, J. P.; Mayeux, P.; Romagnolo, B.; Luquet, S.; Bossard, P.; Perret, C., Lkb1 suppresses amino acid-driven gluconeogenesis in the liver. Nat Commun 2020, 11 (1), 6127.

22. Shaw, R. J.; Lamia, K. A.; Vasquez, D.; Koo, S. H.; Bardeesy, N.; Depinho, R. A.; Montminy, M.; Cantley, L. C., The kinase LKB1 mediates glucose homeostasis in liver and therapeutic effects of metformin. Science 2005, 310 (5754), 1642–6.

23. Patel, K.; Foretz, M.; Marion, A.; Campbell, D. G.; Gourlay, R.; Boudaba, N.; Tournier, E.; Titchenell, P.; Peggie, M.; Deak, M.; Wan, M.; Kaestner, K. H.; Goransson, O.; Viollet, B.; Gray, N. S.; Birnbaum, M. J.; Sutherland, C.; Sakamoto, K., The LKB1-salt-inducible kinase pathway functions as a key gluconeogenic suppressor in the liver. Nat Commun 2014, 5, 4535.

24. Stefkovich, M. L.; Kang, S. W. S.; Porat-Shliom, N., Intravital Microscopy for the Study of Hepatic Glucose Uptake. Curr Protoc 2021, 1 (5), e139.

25. Edgar, R.; Domrachev, M.; Lash, A. E., Gene Expression Omnibus: NCBI gene expression and hybridization array data repository. Nucleic Acids Res 2002, 30 (1), 207–10.

26. Kays, J. K.; Shahda, S.; Stanley, M.; Bell, T. M.; O’Neill, B. H.; Kohli, M. D.; Couch, M. E.; Koniaris, L. G.; Zimmers, T. A., Three cachexia phenotypes and the impact of fat-only loss on survival in FOLFIRINOX therapy for pancreatic cancer. J Cachexia Sarcopenia Muscle 2018, 9 (4), 673–684.

27. Lin, X. L.; Liu, Y. B.; Hu, H. J., Metabolic role of fibroblast growth factor 21 in liver, adipose and nervous system tissues. Biomed Rep 2017, 6 (5), 495–502.

28. Inagaki, T.; Dutchak, P.; Zhao, G.; Ding, X.; Gautron, L.; Parameswara, V.; Li, Y.; Goetz, R.; Mohammadi, M.; Esser, V.; Elmquist, J. K.; Gerard, R. D.; Burgess, S. C.; Hammer, R. E.; Mangelsdorf, D. J.; Kliewer, S. A., Endocrine regulation of the fasting response by PPARalpha-mediated induction of fibroblast growth factor 21. Cell Metab 2007, 5 (6), 415–25.

29. Oost, L. J.; Kustermann, M.; Armani, A.; Blaauw, B.; Romanello, V., Fibroblast growth factor 21 controls mitophagy and muscle mass. J Cachexia Sarcopeni 2019, 10 (3), 630–642.

30. Jung, H. W.; Park, J. H.; Kim, D.; Jang, I. Y.; Park, S. J.; Lee, J. Y.; Lee, S.; Kim, J. H.; Yi, H. S.; Lee, E.; Kim, B. J., Association between serum FGF21 level and sarcopenia in older adults. Bone 2021, 145.

31. Pagliassotti, M. J.; Myers, S. R.; Moore, M. C.; Neal, D. W.; Cherrington, A. D., Magnitude of Negative Arterial-Portal Glucose Gradient Alters Net Hepatic Glucose Balance in Conscious Dogs. Diabetes 1991, 40 (12), 1659–1668.

32. Sokolovic, M.; Sokolovic, A.; Wehkamp, D.; Ver Loren van Themaat, E.; de Waart, D. R.; Gilhuijs-Pederson, L. A.; Nikolsky, Y.; van Kampen, A. H.; Hakvoort, T. B.; Lamers, W. H., The transcriptomic signature of fasting murine liver. BMC Genomics 2008, 9, 528.

33. Dean, E. D.; Li, M. Y.; Prasad, N.; Wisniewski, S. N.; Von Deylen, A.; Spaeth, J.; Maddison, L.; Botros, A.; Sedgeman, L. R.; Bozadjieva, N.; Ilkayeva, O.; Coldren, A.; Poffenberger, G.; Shostak, A.; Semich, M. C.; Aamodt, K. I.; Phillips, N.; Yan, H.; Bernal-Mizrachi, E.; Corbin, J. D.; Vickers, K. C.; Levy, S. E.; Dai, C. H.; Newgard, C.; Gu, W.; Stein, R.; Chen, W. B. A.; Powers, A. C., Interrupted Glucagon Signaling Reveals Hepatic alpha Cell Axis and Role for L-Glutamine in alpha Cell Proliferation. Cell Metabolism 2017, 25 (6), 1362-+.

34. Pei, L.; Waki, H.; Vaitheesvaran, B.; Wilpitz, D. C.; Kurland, I. J.; Tontonoz, P., NR4A orphan nuclear receptors are transcriptional regulators of hepatic glucose metabolism. Nat Med 2006, 12 (9), 1048–55.

35. Min, A. K.; Bae, K. H.; Jung, Y. A.; Choi, Y. K.; Kim, M. J.; Kim, J. H.; Jeon, J. H.; Kim, J. G.; Lee, I. K.; Park, K. G., Orphan nuclear receptor Nur77 mediates fasting-induced hepatic fibroblast growth factor 21 expression. Endocrinology 2014, 155 (8), 2924–31.

36. Settembre, C.; De Cegli, R.; Mansueto, G.; Saha, P. K.; Vetrini, F.; Visvikis, O.; Huynh, T.; Carissimo, A.; Palmer, D.; Klisch, T. J.; Wollenberg, A. C.; Di Bernardo, D.; Chan, L.; Irazoqui, J. E.; Ballabio, A., TFEB controls cellular lipid metabolism through a starvation-induced autoregulatory loop. Nature Cell Biology 2013, 15 (6), 647-+.

37. Samuel, V. T.; Shulman, G. I., Nonalcoholic Fatty Liver Disease as a Nexus of Metabolic and Hepatic Diseases. Cell Metabolism 2018, 27 (1), 22–41.

38. Brooks, S. P. J., Chapter 9 Fasting and refeeding: Models of changes in metabolic efficiency. In Cell and Molecular Response to Stress, Storey, K. B.; Storey, J. M., Eds. Elsevier: 2001; Vol. 2, pp 111–127.

39. Farooq, A.; Chamari, K.; Sayegh, S.; El Akoum, M.; Al-Mohannadi, A. S., Ramadan daily intermittent fasting reduces objectively assessed habitual physical activity among adults. Bmc Public Health 2021, 21 (1).

40. Minghelli, G.; Schutz, Y.; Whitehead, R.; Jequier, E., Seasonal changes in 24-h and basal energy expenditures in rural Gambian men as measured in a respiration chamber. Am J Clin Nutr 1991, 53 (1), 14–20.

41. Berndt, N.; Horger, M. S.; Bulik, S.; Holzhutter, H. G., A multiscale modelling approach to assess the impact of metabolic zonation and microperfusion on the hepatic carbohydrate metabolism. PLoS Comput Biol 2018, 14 (2), e1006005.

42. Halpern, K. B.; Shenhav, R.; Matcovitch-Natan, O.; Toth, B.; Lemze, D.; Golan, M.; Massasa, E. E.; Baydatch, S.; Landen, S.; Moor, A. E.; Brandis, A.; Giladi, A.; Avihail, A. S.; David, E.; Amit, I.; Itzkovitz, S., Single-cell spatial reconstruction reveals global division of labour in the mammalian liver. Nature 2017, 542 (7641), 352–356.

43. Yang, J.; MacDougall, M. L.; McDowell, M. T.; Xi, L.; Wei, R.; Zavadoski, W. J.; Molloy, M. P.; Baker, J. D.; Kuhn, M.; Cabrera, O.; Treadway, J. L., Polyomic profiling reveals significant hepatic metabolic alterations in glucagon-receptor (GCGR) knockout mice: implications on anti-glucagon therapies for diabetes. BMC Genomics 2011, 12, 281.

44. Benhamouche, S.; Decaens, T.; Godard, C.; Chambrey, R.; Rickman, D. S.; Moinard, C.; Vasseur-Cognet, M.; Kuo, C. J.; Kahn, A.; Perret, C.; Colnot, S., Apc tumor suppressor gene is the “zonation-keeper” of mouse liver. Dev Cell 2006, 10 (6), 759–70.

45. Sekine, S.; Lan, B. Y.; Bedolli, M.; Feng, S.; Hebrok, M., Liver-specific loss of beta-catenin blocks glutamine synthesis pathway activity and cytochrome p450 expression in mice. Hepatology 2006, 43 (4), 817–25.

46. Charawi, S.; Just, P. A.; Savall, M.; Abitbol, S.; Traore, M.; Metzger, N.; Ravinger, R.; Cavard, C.; Terris, B.; Perret, C., LKB1 signaling is activated in CTNNB1-mutated HCC and positively regulates beta-catenin-dependent CTNNB1-mutated HCC. J Pathol 2019, 247 (4), 435–443.

47. Spicer, J.; Rayter, S.; Young, N.; Elliott, R.; Ashworth, A.; Smith, D., Regulation of the Wnt signalling component PAR1A by the Peutz-Jeghers syndrome kinase LKB1. Oncogene 2003, 22 (30), 4752–4756.

48. Sun, T. Q.; Lu, B. W.; Feng, J. J.; Reinhard, C.; Jan, Y. N.; Fantl, W. J.; Williams, L. T., PAR-1 is a Dishevelled-associated kinase and a positive regulator of Wnt signalling. Nature Cell Biology 2001, 3 (7), 628–636.

49. Zhan, Y. Y.; Chen, Y.; Zhang, Q.; Zhuang, J. J.; Tian, M.; Chen, H. Z.; Zhang, L. R.; Zhang, H. K.; He, J. P.; Wang, W. J.; Wu, R.; Wang, Y.; Shi, C.; Yang, K.; Li, A. Z.; Xin, Y. Z.; Li, T. Y.; Yang, J. Y.; Zheng, Z. H.; Yu, C. D.; Lin, S. C.; Chang, C.; Huang, P. Q.; Lin, T.; Wu, Q., The orphan nuclear receptor Nur77 regulates LKB1 localization and activates AMPK. Nat Chem Biol 2012, 8 (11), 897–904.

50. Gougelet, A.; Torre, C.; Veber, P.; Sartor, C.; Bachelot, L.; Denechaud, P. D.; Godard, C.; Moldes, M.; Burnol, A. F.; Dubuquoy, C.; Terris, B.; Guillonneau, F.; Ye, T.; Schwarz, M.; Braeuning, A.; Perret, C.; Colnot, S., T-cell factor 4 and beta-catenin chromatin occupancies pattern zonal liver metabolism in mice. Hepatology 2014, 59 (6), 2344–57.

